# siProGenA: Generative siRNA Candidate Construction via Position Proposal and Guide Generation

**DOI:** 10.64898/2026.08.31.748303

**Authors:** Zhiqi Ma, Jiale Zhou, Rubo Wang, Zhipeng Deng, Zhijian Wu, Yefeng Zheng

**Affiliations:** School of Engineering, Westlake University, Hangzhou, China; Shanghai Artificial Intelligence, Laboratory, Shanghai, China

**Keywords:** siRNA candidate construction, candidate-position proposal, guide-sequence generation, discrete diffusion, Bayesian flow networks

## Abstract

Small interfering RNAs (siRNAs) are short guide RNAs that recruit the RNA-induced silencing complex (RISC) to complementary target sites on messenger RNAs (mRNAs), triggering Ago2-mediated cleavage and gene silencing. siRNA design requires compact candidate sets that cover a target while preserving efficacy, specificity, and practical sequence constraints. Existing pipelines usually enumerate candidate windows, assign a canonical guide to each window, and then rank preconstructed siRNA–mRNA pairs. This has produced strong pairwise efficacy predictors, but leaves a candidate-construction gap: candidate positions and guide sequences are fixed before the model begins to rank them. We address this gap by decomposing siRNA candidate construction into two generative decisions: where to place candidates within an mRNA segment, and what constrained guide variants to consider at a candidate position. We instantiate this framework as siProGenA, using a Discrete Denoising Diffusion Probabilistic Model (D3PM) for mRNA-conditioned position proposal and a Bayesian Flow Network (BFN) for temperature-controlled guide generation. On 62 positive test segments, the diversity-aware final library reaches Hit@1 = 0.790 and Hit@5 = 0.903. In a measured-site controlled Stage 2 evaluation, seed- and cleavage-preserving variants outscore the canonical complement for 89.8% of measured sites, with supporting gains across additional computational scorers, random-mismatch controls, and biophysical diagnostics. Together, the results support a modular proposal–generation view of siRNA candidate construction for prioritizing compact candidate sets.

## 1 Introduction

Small interfering RNAs guide the RNA-induced silencing complex (RISC) to target sites on mRNAs, where guide–target pairing enables Ago2-mediated cleavage and gene silencing [7]. In this task, an siRNA guide is a short RNA sequence designed to pair with a target messenger RNA (mRNA) region and induce knockdown of the corresponding transcript. Computational siRNA design has therefore made steady progress as a pairwise efficacy prediction problem: scan the mRNA with candidate target windows, assign a guide to each window, score each candidate independently, and rank or filter the resulting list [3, 21, 28]. This scoring paradigm is useful and biologically grounded, but it leaves the construction of the candidate set largely outside the learning problem. We refer to this limitation as the *candidate-construction gap*: the model is asked to rank candidates after the candidate positions and guide sequences have already been fixed.

The candidate-construction gap hides two design decisions that matter in practice. The first is **where to place candidates**. A design process must choose candidate positions from a longer mRNA segment, yet adjacent 19-mer windows share most of their nucleotides and often occupy the same local sequence context. Independent ranking can therefore waste a limited candidate budget on redundant nearby windows instead of proposing a diverse set of promising sites. The second is **which guide sequence to consider once a candidate position is identified**. The Watson–Crick reverse complement remains the canonical guide candidate, but it defines only one candidate around that anchor. Constrained variants around the same candidate position may expose additional viable guide candidates. This is biologically plausible because guide–target pairing is position-dependent, and selected mismatches outside the seed and central cleavage-sensitive positions can be tolerated [10, 12, 17]. These two decisions define the candidate pool; practical selection can then impose spacing, seed-uniqueness, toxicity, and off-target constraints.

To close this gap, we move siRNA design from post-hoc ranking of fixed candidates to generative candidate construction. We propose siProGenA, which decomposes the problem into two modeling stages and one selection step. Stage 1 performs mRNA-conditioned candidate position proposal: given a reconstructed target segment, it models a distribution over candidate cleavage sites rather than scoring each window in isolation, and a diversity-aware reranker converts this dense proposal signal into a compact top-*k* candidate set. Stage 2 performs position-conditioned guide-sequence generation: given a candidate position and the canonical reverse complement as the anchor candidate, it generates constrained guide variants that preserve seed and cleavage-site complementarity while allowing controlled variation elsewhere. In deployment, Stage 2 can be applied to positions proposed by Stage 1; in our controlled Stage 2 evaluation, we isolate guide generation by conditioning on experimentally measured test sites.

We instantiate Stage 1 with a Discrete Denoising Diffusion Probabilistic Model (D3PM) [2] over ordered target positions, and Stage 2 with a Bayesian Flow Network (BFN) [8] over 19-mer guide sequences. Stage 2 generates constrained guide variants around the canonical complement, with Best-of-*N* distillation used to bias generation toward computationally favorable variants.

Experiments evaluate both design decisions. On 62 target segments with experimentally validated active sites, Stage 1 achieves final-library Hit@5 = 0.903, outperforming per-site predictors at the compact-library level. For Stage 2, seed-cleavage-mid constrained variants outscore the canonical complement for 89.8% of measured sites at *T* = 1.0 in a measured-site controlled evaluation. Supporting analyses show consistent gains across computational scorers, difficulty strata, random mismatch controls, GC-content checks, and ViennaRNA diagnostics.

Our contributions are:

- We identify the candidate-construction gap in enumerate– score–rank siRNA pipelines and reformulate siRNA design as two explicit decisions: where to place candidates within an mRNA segment and what guide variants to consider at a candidate position.
- We propose siProGenA, an instantiation that uses D3PM for mRNA-conditioned position proposal, diversity-aware reranking for set construction, and BFN for temperature-controlled guide generation around a candidate position.
- Extensive evaluations show that Stage 1 improves library-level site coverage, while a measured-site controlled Stage 2 evaluation shows that constrained guide variants outperform the canonical complement across external scorers, composition checks, random-mismatch controls, and biophysical diagnostics.

## 2 Related Work

Prior work provides strong scoring models and biological design constraints, but candidate construction is usually handled outside the learned model. We therefore discuss related work in terms of candidate scoring, constrained guide variation, and generative construction.

### Pairwise siRNA efficacy prediction and site selection

Early siRNA design relied on empirical rules [27, 30]; later methods used hand-crafted features [15, 18, 23, 25, 32]; while recent models use Transformers, graph networks, RNA language models, and structure-aware representations [3, 5, 6, 21, 28]. These methods have improved candidate-level efficacy prediction by modeling siRNA sequence features, mRNA context, thermodynamics, and siRNA–mRNA interactions. In design pipelines, however, candidate positions are usually obtained by sliding-window enumeration and then scored or filtered independently. For example, OligoFormer enumerates 19-nt windows across the target mRNA with flanking context, then scores and ranks candidates by predicted efficacy [3]. Thus, existing predictors are strong candidate scorers, while candidate-position proposal remains largely an enumerate-then-rank procedure.

### Guide–target pairing and constrained guide variation

Canonical siRNA design is anchored by Watson–Crick pairing between the guide strand and its target. At the same time, Argonaute-mediated recognition is position-dependent: seed pairing controls target recognition and off-target effects, while central positions are tied to cleavage [4, 10–12, 16, 17]. This motivates a conservative design space for guide variation: complementarity should be preserved at sensitive positions, while variation outside those positions must be controlled rather than unconstrained.

### Generative candidate construction

Discrete diffusion [2, 13] and Bayesian Flow Networks [8] have been used for molecular, protein, and peptide generation [1, 9, 29, 33]. Recent diffusion-based siRNA work generates siRNA–mRNA pairs directly [24]. These generative models demonstrate the utility of discrete sequence generation, but they do not directly address compact siRNA library construction from mRNA-conditioned site proposal followed by constrained guide variation.

## 3 Problem Definition

Let x = (*x*_1_, …, *x*_*L*_)be a target segment over the RNA alphabet {*A, C, G, U* }of length *L*. A candidate siRNA is defined by its guide-strand start index *i ∈* {1, …, *L −* 18 }and the 19-nucleotide target window t_*i*_ = (*x*_*i*_, … *x*_*i* +18_). We follow the standard 19-nt guide setting used by the source assays and prior siRNA efficacy predictors. The complementary guide strand 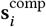 = revcomp (t_*i*_) is determined by Watson–Crick base pairing and serves as the default design for each site. More generally, each site has an admissible guide set *S*_*i*_ *⊂* {*A, C, G, U* }^19^ that contains 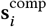 and variants that preserve complementarity at protected positions such as the seed and cleavage-sensitive sites. Depending on the constraint mode, additional positions, including the 3^′^ supplementary region, may also be protected. Stage 2, described in Section 4.2, samples candidate guide variants from *S*_*i*_ to expand the candidate set at selected sites.

For each target segment we have experimentally measured data on a subset of candidate sites: a binary label *y*_*i*_ *∈* {0, 1} indicating *≥*70% knockdown, and a continuous efficacy score *e*_*i*_ *∈* [0, 1]. Let *A* (x) = {*i*: *y*_*i*_ = 1} denote the validated active sites.

The final output is a set of *k* candidates {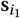, …, 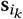}, where each site *i*_*j*_ is assigned a guide strand. We decompose this objective into two modeling problems: identify promising candidate positions on the mRNA segment and generate admissible guide variants for selected positions. The assembled library should cover *A*(x) while satisfying safety and diversity constraints.

## 4 siProGenA Framework

The framework closes the candidate-construction gap by making the two hidden design decisions explicit. Stage 1 addresses *where to place candidates*: it proposes candidate positions across the reconstructed mRNA segment instead of ranking each measured window in isolation. Stage 2 addresses *what guide variants to consider*: it expands a candidate position around the canonical reverse complement under biological constraints. The final selection step turns the constructed candidates into a practical compact set by applying sequence, safety, and diversity constraints.

This decomposition also matches the geometry of the two output spaces. Site proposal operates on a small ordered categorical space, the *N* = *L −*18 possible cleavage positions. D3PM with structured transitions [2] defines a categorical diffusion over these indices, allowing sparse experimental labels to propagate through a segment-level encoder. Guide generation operates on a 4^19^sequence space. BFN treats each guide position as a continuous-time Bayesian update, refines all 19 positions jointly, and exposes temperature as an inference-time control for diversity. The ablations in Section 5.3 validate this space-specific pairing.

### 4.1 Stage 1: mRNA-Conditioned Position Proposal

#### Module role

Stage 1 addresses the where-to-place-candidates side of the candidate-construction gap. Given a target segment x, it returns a distribution over the *N* = *L −* 18 candidate cleavage positions. This differs from enumerate-then-rank selection: nearby 19-mer windows compete for probability mass before the final candidate budget is spent.

The Stage 1 output is a distribution over an ordered set of candidate target positions, not an independent label for each 19-mer. This formulation is useful because neighboring windows share most nucleotides and only a sparse subset of positions is experimentally measured. We therefore model position proposal as denoising a corrupted categorical distribution over positions. The forward process gradually mixes the measured activity distribution with a uniform position prior, and the reverse model learns to recover validated active positions from the encoded mRNA segment. This multi-step formulation gives the model a structured way to refine uncertain position proposals, which we later isolate against single-step direct heads in Section 5.3.

#### Target segment encoder

The input x is one-hot encoded with four nucleotide channels and one additional channel for unknown bases, then linearly projected to *d*_model_ = 128 dimensions with sine-cosine positional encodings [31]. A 4-layer Transformer encoder with 4 attention heads and feed-forward layers with dimension 512 processes the sequence into per-position representations H *∈* ℝ^*L*×128^. These representations are pooled into site embeddings 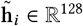 by combining the mean over each 19-mer window (*x*_*i*_, …, *x*_*i*+18_) with the center-position representation:

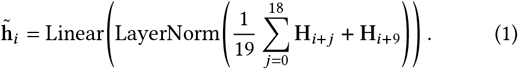

#### Discrete diffusion over candidate positions

Let *N* = *L−*18 be the number of candidates. We treat the problem as a categorical diffusion over these *N* positions rather than over nucleotide identities. The D3PM state space is the set of candidate site indices, and each position’s probability mass represents its suitability as a cleavage site. The ground-truth target is a *soft distribution π*_0_ *∈* Δ^*N −*1^: for a target segment with measured sites, the probability mass is distributed over measured positions using a temperature-scaled softmax of efficacy scores, and zero elsewhere. Target segments with no measured sites use a uniform baseline over valid positions. At diffusion step *t∈* { 1, …, *T*} (*T* = 100), the forward transition matrix Q_*t*_*∈* ℝ^*N* ×*N*^ applies uniform-absorbing corruption toward a uniform distribution over the *N* positions:

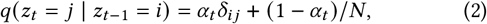

where the per-step α_*t*_ is chosen so that the cumulative retention factor follows a cosine schedule with offset *s* = 0.008:

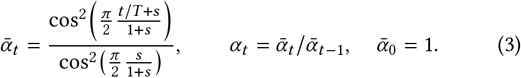

For a soft target distribution, the marginal corrupted distribution is tractable:

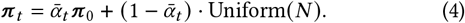

The reverse process is parameterized as a direct *π*_0_ prediction. The PosHead takes site embeddings 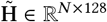, a corrupted sampled index *z*_*t*_ *~ π*_*t*_, and a sinusoidal time embedding, and produces logits *ℓ*_*i*_ for each position via a multilayer perceptron (MLP) with feature-wise linear modulation (FiLM) conditioning [26] and a learned bias at the *z*_*t*_ position. Let 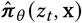 = softmax (*ℓ*) denote the predicted clean-site distribution. The training loss is the cross-entropy between this predicted distribution and the soft target:

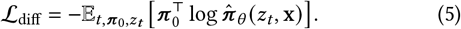

#### Training objective

At training time, a diffusion step *t ~ U* (1, *T*) is sampled uniformly, the corrupted state *z*_*t*_ is sampled from the marginal distribution 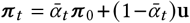, where u is the uniform distribution over positions. `PosHead` predicts 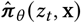 by applying a softmax over the *N* position logits. The structured uniform-absorbing transition makes this marginal tractable in closed form, so each training step requires only a single forward pass. The model produces logits over valid candidate positions and handles different segment lengths through the position mask. Unmeasured sites receive near-zero mass in *π*_0_, and the diffusion process propagates information from sparse labels through the target segment-level encoder.

#### Auxiliary supervision

Two additional heads share the site embeddings: an efficacy regression head, denoted `eff_head`, trained with smooth L1 loss, and an activity-classification head, denoted `cls_head`, trained with cross-entropy loss to predict *≥*70% knock-down. A pairwise ranking loss contrasts the predicted efficacy differences between randomly sampled site pairs with moderate measured efficacy differences. These auxiliary objectives stabilize training when the diffusion signal is noisy at early steps and are also reused as scoring components during constrained selection, described in Section 4.3. The total loss is *ℒ*_1_ = *ℒ*_diff_ +*λ*_eff_ *ℒ*_eff_ + *λ*_cls_ *ℒ*_cls_ + *λ*_pair_ *ℒ*_pair_, with *λ*_eff_ = 0.5, *λ*_cls_ = 0.5, *λ*_pair_ = 2.0.

During inference, we initialize from the uniform stationary distribution and run*T* reverse steps to obtain site logits over {1, …, *N*}. The site-level score used for Stage 1 reranking combines the raw D3PM position logit with the auxiliary activity-classification probability and efficacy prediction. A greedy diversity-aware decoder then selects a nonredundant top-*k* set by adding a normalized Hamming distance bonus to the candidate’s closest already selected site. We compare direct ancestral sampling, diversity-aware reranking, and deterministic ranking with the efficacy or activity-classification head in Section 5.3 to attribute the final library gain to proposal quality rather than only to post-hoc set conversion.

### 4.2 Stage 2: Position-Conditioned Guide Generation

#### Module role

Stage 2 addresses the what-guide-to-use side of the candidate-construction gap. Given a candidate position *i* and its canonical reverse-complement guide 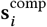, it generates admissible 19-mer guide variants s_*i*_*∈ S*_*i*_. The goal is candidate expansion around a proposed or measured position, not replacement of the Stage 1 position proposal. For deployment, the candidate position can come from Stage 1. For the controlled Stage 2 experiments in Section 5.4, we instead condition on experimentally measured test sites to evaluate guide generation separately from position proposal.

This is different from scoring a fixed siRNA–mRNA pair. The relevant properties, including GC content, seed preservation, mismatch placement, and duplex stability, are properties of the full guide sequence. BFN is a natural fit because it refines all 19 guide positions jointly and uses temperature as a training-free control over the diversity–efficacy trade-off.

#### BFN generation process

Bayesian Flow Networks model guide generation as iterative refinement of a categorical belief over the 19 guide positions. We use a communication protocol over *T*_bfn_ refinement steps with a noise schedule *β t* = *β*_max_ (*t T*_bfn_)^2^, with *β*_max_ = 1.0. For a true discrete sequence s over *K* = 4 nucleotides, the sender emits a noisy observation y | s *~* N (*β* (*K*e_s_*−*1), *β K*I), where e_s_ is the one-hot encoding. The receiver maintains a Bayesian posterior belief *θ*_*t*_ *∈* (Δ^*K −*1^)^19^ over each position’s categorical distribution. It is initialized as uniform and updated by Bayes rule, *θ*_*t*_ ∝ *θ*_*t* 1_ exp r_*t*_, where r_*t*_ denotes the noisy message received at refinement step *t*. A 6-layer Transformer backbone maps the current belief to *p*_rec_ 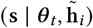 using cross-attention to the Stage 1 site embedding. The training goal is to minimize the variational bound

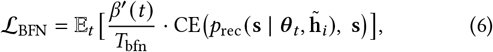

which weights the reconstruction cross-entropy by the instantaneous precision rate *β*^′^(*t*) / *T*_bfn_. During inference, temperature controlled sampling is achieved by scaling the receiver’s predicted logits *ℓ* before the softmax:

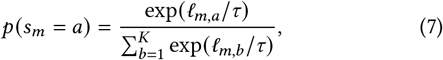

where *τ >* 1 flattens the distribution and *τ* = 1 recovers the learned softmax distribution, with *τ*→ 0 approaching argmax sampling.

This is a zero-cost, inference-only intervention that smoothly interpolates between diverse and concentrated generation.

Because BFN refines all 19 positions jointly, it captures global sequence properties that left-to-right models miss. Site-level guidance and sequence regularization are provided separately through the Best-of-*N* distillation procedure below.

#### Training via Best-of-*N* distillation

Because training data contain only one measured siRNA per site, we use Best-of-*N* distillation as a practical training procedure: the model samples multiple constrained guide variants, scores them with a composite computational proxy, and fine-tunes on the top-scoring variants. This encourages admissible high-scoring variants without requiring multiple experimentally measured guides per site. The proxy combines guide-level efficacy scores, GC-range regularization, and hard constraint violation penalties.

#### Sequence-level constrained sampling

Unconstrained sampling can corrupt seed and cleavage-site complementarity, producing biologically implausible variants. To ensure validity, we constrain sampling at inference time. During each Bayesian refinement step, the belief update for target positions corresponding to the seed and cleavage site is clamped to the complementary nucleotide, allowing mismatches elsewhere in the 19-mer, including the 5^′^ terminus and the 3^′^ supplementary and terminus regions, where RISC-mediated target recognition is more tolerant [10, 12, 17]. Clamping replaces the model-predicted distribution at protected positions with a point mass on the complementary base, so the generated guide preserves the protected pairing pattern by construction.

Intuitively, BFN treats generation as gradually refining a belief over each position: at early steps the model sees nearly uniform distributions and makes coarse predictions; at late steps it sees peaked distributions and makes fine-grained predictions. The noise schedule controls how quickly this refinement happens, and the temperature parameter in Equation 7 sharpens or flattens the final sampling distribution without altering the learned belief-update process.

### 4.3 Diversity-Aware Reranking and Constrained Selection

The final output is a compact library rather than a single ranked position. We therefore use two selection steps. The first step reranks Stage 1 position proposals into a nonredundant top-*k* set. The second step, after Stage 2 has generated guide variants, assembles the final constrained library with safety and diversity terms.

**Stage 1 position reranking**. For a guide s, let seed (s) = *s*_2:7_ denote the 6-mer at guide positions 2–7, which is central to seed-mediated off-target recognition [4, 16]. Given a seed-toxicity table with cell viability *ν* (seed) and range [*ν*_min_, *ν*_max_], we define

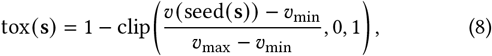

so lower cell viability gives higher toxicity. For off-target risk, let

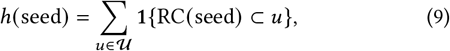

where *U* is the human untranslated region (UTR) reference set.

The normalized off-target proxy is

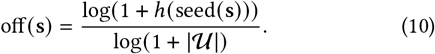

For any candidate guide g and a partially selected set ℬ, we use one normalized Hamming diversity term:

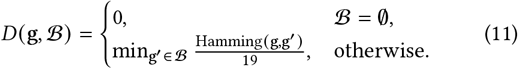

For each candidate position *i*, Stage 1 produces a D3PM position logit 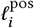, an activity-classification probability 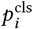, and an efficacy prediction *e*_*i*_. Let c_*i*_ be the canonical complementary guide for position *i*. The per-position score is

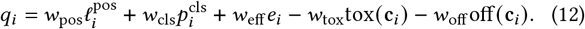

Here 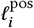 is the raw PosHead logit, not a calibrated softmax probability. Because the PosHead is trained against an efficacy-weighted soft target distribution, this logit is best interpreted as a relative position-proposal signal.

Reranking is greedy. Let *S* be the set of already selected sites and *C*_*S*_ = { c_*j*_: *j ∈ S* } be their canonical guides. At each step we select

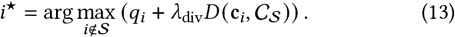

This layer uses proposal and auxiliary efficacy signals plus sequence diversity, toxicity, and off-target proxies. The weights are selected on the validation set and fixed before test evaluation.

#### Constrained library assembly

After Stage 2 generates guide variants, each candidate *a* = (*i*, s) inherits the Stage 1 site signals and receives guide-specific toxicity, off-target, and sequence-diversity scores. During final selection, the candidate score is

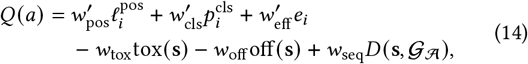

where *A* is the partially assembled library and *G*_*A*_ = {s_*j*_: (*j*, s_*j*_) *∈* A}. The assembly weights are selected on the validation set and fixed before test evaluation.

## 5 Experiments

The experiments are organized around three research questions (R*Q*s). RQ1 asks whether segment-level position proposal improves library-level coverage over per-site ranking. R*Q*2 identifies which Stage 1 components account for the gain. RQ3 tests whether position-conditioned guide generation can improve over the canonical complementary guide under external evaluators. We also report supporting diagnostics for Stage 2 to check composition, difficulty dependence, and random mismatch controls.

### 5.1 Setup

#### Dataset

We use the Huesken [14], Mix, and Takayuki sources compiled by OligoFormer [3]. Each record contains a 19-nt siRNA guide and a 57-nt mRNA target window from reporter assays. To avoid overlap of near-identical window contexts across splits, we first group records by transcript/target identity, merge overlapping windows within each transcript into reconstructed target segments, and assign all segments from the same transcript to the same train, validation, or test split. The resulting dataset contains 1,258 reconstructed segments, including 726 with at least one validated active site (*≥* 70% knockdown). We use an approximately 80/10/10 transcript-level split; the test split contains 120 segments, including 62 positive segments.

#### Evaluation metrics

For position proposal, we report **Hit@K**, the fraction of test target segments whose selected library contains at least one active site, at budget *k* = 1 and *k* = 5. We also report Recall@5, the fraction of measured active sites recovered among the top five predictions for each segment, and MRR@5, the mean reciprocal rank of the first recovered active site within the top-five selected library, with zero assigned when no active site is recovered. For guide generation, we evaluate measured-site variants with external computational scorers and report predicted efficacy, gen*>*comp win rate, GC content (the fraction of G/C nucleotides), mismatch counts, random-mismatch controls, and ViennaRNA binding diagnostics.

#### Baselines

We compare against: OligoFormer [3], siRNABERT [34], siRNADiscovery [21], DSIR [32], siRNAPred [18], OligoWalk [23], s-BioPredsi [19], and Monopoli-RF [25]. Learned baselines are retrained or calibrated on the same transcript-level train/validation/test split when training data are available. All baselines produce persite scores; we select top-*k* positions for library-level evaluation. The main text reports representative learned and traditional baselines, with an expanded baseline list in Appendix A.1.

#### Implementation

Stage 1 uses a 4-layer Transformer and a D3PM with *T* = 100, uniform-absorbing transitions, and a cosine 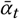 schedule. It is trained for 50 epochs with AdamW, batch size 4, 4-step gradient accumulation, and 200 warmup steps. Stage 2 uses a 6-layer BFN Transformer trained via Best-of-*N* distillation and evaluated with a pre-trained OligoFormer pairwise evaluator. The model is trained on a single RTX 4080 SUPER and takes approximately 6 hours for Stage 1 and 4 hours for Stage 2.

### 5.2 RQ1: Segment-Level Target-Site Proposal

Table 1 answers RQ1 by comparing segment-level proposal with top-*k* libraries derived from representative per-site scoring base-lines. siProGenA reports the full Stage 1 output: D3PM position proposal followed by diversity-aware reranking from Section 4.3. The final compact library outperforms the strongest representative top-*k* baselines. The result suggests that the proposal scores and the subsequent reranking step provide a more useful selected set for compact library construction

**Takeaway. Stage 1 improves library-level top-***k* **recovery over representative per-site baselines, with reranking converting proposal scores into a compact library**.

**Table 1:** Stage 1 comparison against the strongest representative baselines on the test set. All methods are converted into libraries by selecting top-*k* positions; an expanded baseline list is reported in Appendix A.1.

| Method | Hit@5 | Recall@5 | MRR@5 |
| --- | --- | --- | --- |
| Sliding window uniform | 0.597 | 0.493 | 0.373 |
| DSIR [32] | 0.387 | 0.264 | 0.279 |
| siRNABERT [34] | 0.355 | 0.265 | 0.190 |
| siRNAPred [18] | 0.355 | 0.283 | 0.217 |
| OligoFormer [3] | 0.323 | 0.272 | 0.171 |
| siProGENA final library | **0.903** | **0.813** | **0.840** |

The strongest non-neural coverage baseline is sliding-window uniform, which is stronger than several learned scorers but still falls well below siProGenA on Hit@5 and Recall@5. An expanded baseline list is deferred to Appendix A.1.

### 5.3 RQ2: Attribution of the Stage 1 Gain

RQ2 asks where the Stage 1 gain comes from. Tables 2–4 report the mechanism-level ablations used to identify which components drive the library-level gain: the multi-step proposal head, the segment encoder, auxiliary heads, and post-hoc reranking.

**Table 2:**
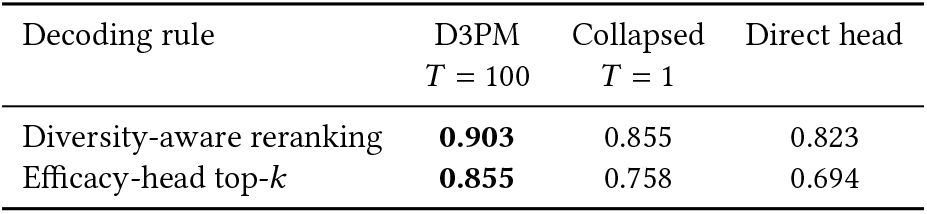
Stage 1 mechanism ablation (Hit@5). “Diversity-aware reranking” selects a nonredundant top-5 library from proposal scores, while “efficacy-head top-*k*” ranks positions by the auxiliary efficacy head. The pure direct head keeps the same encoder and reranking protocol but replaces D3PM with a single linear position head.

**Takeaway. The gain comes mainly from the multi-step D3PM proposal distribution: single-step heads and alternative schedules reduce coverage, while reranking turns dense proposal scores into a compact nonredundant library**.

**Diffusion vs. direct classifier**. We test whether the multi-step diffusion head is necessary, or whether a single-step classifier over position embeddings could produce similar results. We compare D3PM with *T* = 100 against two ablations in Table 2: a collapsed diffusion with *T* = 1 that retains the diffusion machinery and a pure feed-forward linear head with no time embedding or diffusion schedule. We compare two decoding rules: diversity-aware reranking selects a nonredundant library from proposal scores, while efficacy head top-*k* ranks sites directly by predicted efficacy. D3PM with diversity-aware reranking exceeds either single-step alternative. The observation is consistent across decoding strategies: iterative diffusion refinement adds position proposal quality beyond a direct classifier over the same segment representation.

The key control is “pure direct head + diversity-aware reranking”: it uses the same segment-level Transformer encoder and the same diversity-aware reranking as full D3PM, but removes the multi-step proposal process. The remaining gap therefore isolates the diffusion proposal mechanism from input context and reranking. Read column-wise, Table 2 shows the same pattern within each decoding strategy, making the attribution stronger than a single headline comparison.

#### Diffusion schedule

Table 3 asks whether the choice of generative schedule matters for the raw ordered proposal distribution before diversity-aware set reranking. Discrete Flow Matching (DFM) [20] is competitive, but D3PM is the most stable schedule on both Hit@1 and Hit@5; BFN trails on this position-proposal task. We therefore use D3PM for Stage 1 and reserve BFN for the 19-mer sequence space, where its temperature control is more useful.

**Table 3:** Stage 1 raw proposal schedule ablation before diversity-aware reranking.

| Schedule | Raw Hit@1 | Raw Hit@5 |
| --- | --- | --- |
| D3PM | **0.758** | **0.887** |
| DFM | 0.742 | 0.887 |
| BFN | 0.726 | 0.774 |

#### Head ablation

Table 4 evaluates the roles of the diffusion proposal, classification, and efficacy heads. Removing the diffusion proposal head collapses coverage, showing that the main Stage 1 signal is the proposal distribution itself. Removing the efficacy head leaves Hit@5 unchanged but lowers MRR@5, suggesting that the efficacy signal is not required for top-5 coverage but helps preserve ranking quality. We retain the efficacy head for its auxiliary role: it provides a dense regression signal during training and a continuous candidate-level score reused during constrained assembly. Removing the activity-classification head, in contrast, raises Hit@5 and Recall@5 but severely degrades reciprocal-rank diagnostics. The full model is therefore a practical multi-objective setting rather than the single best configuration for each diagnostic metric.

**Table 4:** Stage 1 head ablation on the test set. “Full model” uses the diffusion proposal, activity-classification head, and efficacy head. Each ablation removes one component while keeping the remaining training and selection protocol unchanged.

| Stage 1 variant | Hit@5 | Recall@5 | MRR@5 |
| --- | --- | --- | --- |
| Full model | 0.903 | 0.813 | **0.840** |
| w/o activity head | **0.952** | **0.865** | 0.268 |
| w/o efficacy head | 0.903 | 0.846 | 0.564 |
| w/o proposal head | 0.194 | 0.127 | 0.108 |

#### Diversity-aware reranking effect

The main siProGenA library result includes the diversity-aware reranking layer in Section 4.3. Reranking turns a strong dense proposal distribution into the actual library object by selecting a nonredundant top-*k* set, supporting the intended Stage 1 decomposition: D3PM estimates where activity is likely, and diversity-aware reranking converts that signal into a usable candidate library. Appendix A.2 reports a sensitivity analysis showing stable coverage across representative coverage–safety–diversity trade-offs.

### 5.4 RQ3: Position-Conditioned Guide Generation

Stage 2 is evaluated as a controlled guide-variant generation task. To isolate guide generation from position proposal, we condition the generator on experimentally measured candidate positions in the test set and compare each generated variant with the canonical complementary guide at the same position. This evaluation does not use Hit@K, which is a site-coverage metric for Stage 1; instead, it uses external efficacy predictors, gen*>*comp win rate, GC content, mismatch counts, and ViennaRNA binding diagnostics.

The evaluation asks whether BFN can improve the guide around a measured target site while preserving biologically motivated positional constraints, and whether the effect persists under external siRNA scoring models and biophysical diagnostics.

**Takeaway. Seed-cleavage-mid constrained variants improve external predictor scores at both temperatures, with stronger gains at** *T* = 1.0 **and more conservative gains at** *T* = 2.5.

Table 5 asks whether this gain survives biologically motivated positional constraints. It does: the lower-temperature *T* = 1.0 setting improves most sites across all three constraint modes, with the seed-cleavage-mid mode giving the strongest overall trade-off. The full scatter plot and higher-temperature sensitivity check are provided in Appendix A.3.

**Table 5:** Constrained guide generation at the lower-temperature setting *T* = 1.0. OligoFormer scores compare the generated guide with the canonical complement at the same site. VRNA values are changes from the complementary baseline; smaller Δduplex and Δseed indicate better preservation of guide–target binding.

| Constraint mode | OligoFormer orig→gen | $\Delta$ | Win% | GC | Mismatch | VRNA Δduplex / Δseed |
| --- | --- | --- | --- | --- | --- | --- |
| Seed+cleavage | 0.537→0.631 | +0.094 | 82.8% | 0.524 | 8.17 | +20.75 / +2.25 |
| Seed+cleav+mid | **0.537→0.635** | **+0.098** | **89.8%** | 0.484 | 5.20 | +14.72 / +2.24 |
| Only 3' variable | 0.538→0.618 | +0.081 | 89.0% | 0.490 | 3.03 | +6.56 / +1.29 |

Table 6 addresses the main evaluator-bias concern. The same seed-cleavage-mid variants improve mean scores not only under OligoFormer but also under DSIR, s-BioPredsi, and siRNAPred. At *T* = 1.0, all four predictors show strong gains; at *T* = 2.5, the gains are more conservative but remain positive under every predictor. This consensus supports the interpretation that Stage 2 learns useful constrained guide variation rather than exploiting a single scorer.

**Table 6:**
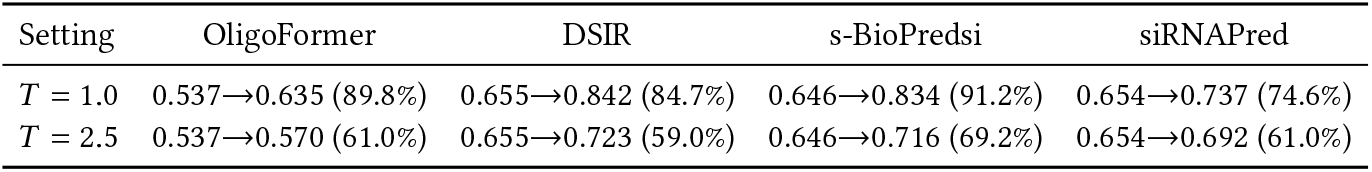
Multi-model consensus for seed-cleavage-mid variants on 354 measured test sites. Each cell reports mean original score→generated score and win rate over the canonical complement.

**Table 7:** Difficulty-stratified Stage 2 gains for the seedcleavage-mid constraint. Sites are grouped by the original complementary guide’s OligoFormer score.

| Temp. | Group | $n$ | Orig | Gen | $\Delta$ | Win rate% |
| --- | --- | --- | --- | --- | --- | --- |
| 1.0 | Hard | 118 | 0.408 | 0.551 | +0.143 | 97.5% |
| 1.0 | Medium | 118 | 0.526 | 0.627 | +0.101 | 91.5% |
| 1.0 | Easy | 118 | 0.677 | 0.726 | +0.049 | 80.5% |
| 2.5 | Hard | 118 | 0.409 | 0.498 | +0.089 | 83.1% |
| 2.5 | Medium | 118 | 0.526 | 0.580 | +0.054 | 69.5% |
| 2.5 | Easy | 118 | 0.677 | 0.634 | -0.043 | 30.5% |

Full temperature and ViennaRNA diagnostics are reported in Appendix A.3 and Appendix A.4; the key point is that the strongest guide generation setting improves predicted efficacy, while constrained variants remain compatible with seed-preserving design.

### 5.5 Supporting Diagnostics for Stage 2

These diagnostics sharpen the interpretation of the Stage 2 result in Section 5.4. They show where guide generation helps, whether the improvement is stronger than a random edit baseline, and whether the variants remain within simple sequence and biophysical design checks.

**Stage 2 gains are strongest where candidate expansion is most needed and are not reproduced by random edits with the same mismatch budget**.

#### Difficulty-stratified gains

We stratify measured sites by the OligoFormer score of the canonical complement. Under the lower-temperature seed-cleavage-mid setting, the largest gain occurs on hard sites. The gain decreases as the canonical guide becomes stronger, which is the desired behavior for a candidate-expansion module: it is most useful where the default complement is weak.

#### Random-mismatch control

Random variants remain near chance under DSIR, s-BioPredsi, and siRNAPred despite using the same edit budget. Table 8 shows that BFN variants are consistently preferred by external scorers while having comparable mismatch, GC, and ViennaRNA perturbation levels. This comparison distinguishes learned guide variation from arbitrary non-complementarity.

**Table 8:** BFN variants versus random mismatches under the same seed-cleavage-mid constraint and mismatch budget at.

| Metric | BFN | Random mismatch |
| --- | --- | --- |
| Mismatch | 5.20 | 5.20 |
| DSIR score / win rate | 0.842 / 84.7% | 0.664 / 49.2% |
| s-BioPredsi score / win rate | 0.834 / 91.2% | 0.667 / 54.0% |
| siRNAPred score / win rate | 0.737 / 74.6% | 0.670 / 53.1% |
| VRNA Δduplex / Δseed | +14.72 / +2.24 | +14.93 / +1.84 |
| GC | 0.484 | 0.537 |

## 6 Conclusion

This work reframes siRNA design from ranking a preconstructed candidate list to constructing the candidate set itself. siProGenA implements this view with mRNA-conditioned position proposal, position-conditioned guide generation, and constrained library selection. Across position-proposal, attribution, and guide-generation experiments, the results show that the main gain comes from proposing candidate positions at the segment level, while controlled guide generation can improve evaluator-predicted efficacy without leaving the constrained guide space.

The same proposal–generation–selection pattern may also be useful for other sequence design tasks where the output is a compact constrained set rather than a single top-ranked candidate.

## 7 Limitations and Ethical Considerations

siProGenA is intended for therapeutic siRNA research and preclinical candidate prioritization; prospective experimental validation and independent checks for efficacy, specificity, and toxicity are required before any biological or clinical use. The data used in this work come from published, de-identified experimental assays and do not involve human-subject data.

## 8 Generative AI Usage

Generative AI tools were used to assist with language editing and organization of the manuscript.

## A Supplementary Material

### A.1 Stage 1 Baselines and Expanded Metrics

The Stage 1 comparison includes neural, rule-based, and thermody-namic baselines. OligoFormer [3], siRNABERT [34], siRNADiscovery [21], siRNAPred [18], and s-BioPredsi [19] are learned siRNA efficacy predictors. DSIR [32] is a rule-based scoring method, while OligoWalk [23] uses thermodynamic accessibility and hybridization calculations. Monopoli-RF [25] is a random-forest predictor trained for siRNA efficacy. All per-site scores are converted into library-level predictions by selecting top-*k* candidate positions. Table 9 reports additional library-level diagnostics at *k* = 5 on the 62 positive test segments.

**Table 9:** Expanded Stage 1 baseline metrics. Hit@5 measures whether at least one validated active site appears in the selected library; Recall@5 measures the fraction of validated active sites recovered; MRR@5 is the mean reciprocal rank of the first recovered active site within the top-five selected library, with zero assigned when no active site is recovered.

| Method | Hit@5 | Recall@5 | MRR@5 |
| --- | --- | --- | --- |
| siProGENA final library | **0.903** | **0.813** | **0.840** |
| Sliding window uniform | 0.597 | 0.493 | 0.373 |
| DSIR-style [32] | 0.387 | 0.264 | 0.279 |
| siRNABERT [34] | 0.355 | 0.265 | 0.190 |
| siRNAPred [18] | 0.355 | 0.283 | 0.217 |
| OligoFormer [3] | 0.323 | 0.272 | 0.171 |
| OligoWalk [23] | 0.323 | 0.248 | 0.141 |
| siRNADiscovery [21] | 0.290 | 0.202 | 0.223 |
| OligoFormer diversity-aware reranking [3] | 0.274 | 0.224 | 0.158 |
| s-BioPredsi [19] | 0.274 | 0.208 | 0.122 |

### A.2 Reranking Weight Sensitivity

Figure 2 evaluates representative coverage–safety–diversity trade-offs in the final reranking layer. The goal is not to optimize weights on the test set, but to check whether the selected library depends on a narrow reranking configuration. Here, None removes safety penalties and diversity rewards; Safety and Diversity enable only the corresponding terms; High div. and High safety increase the corresponding weight relative to the default setting.

**Figure 1.**
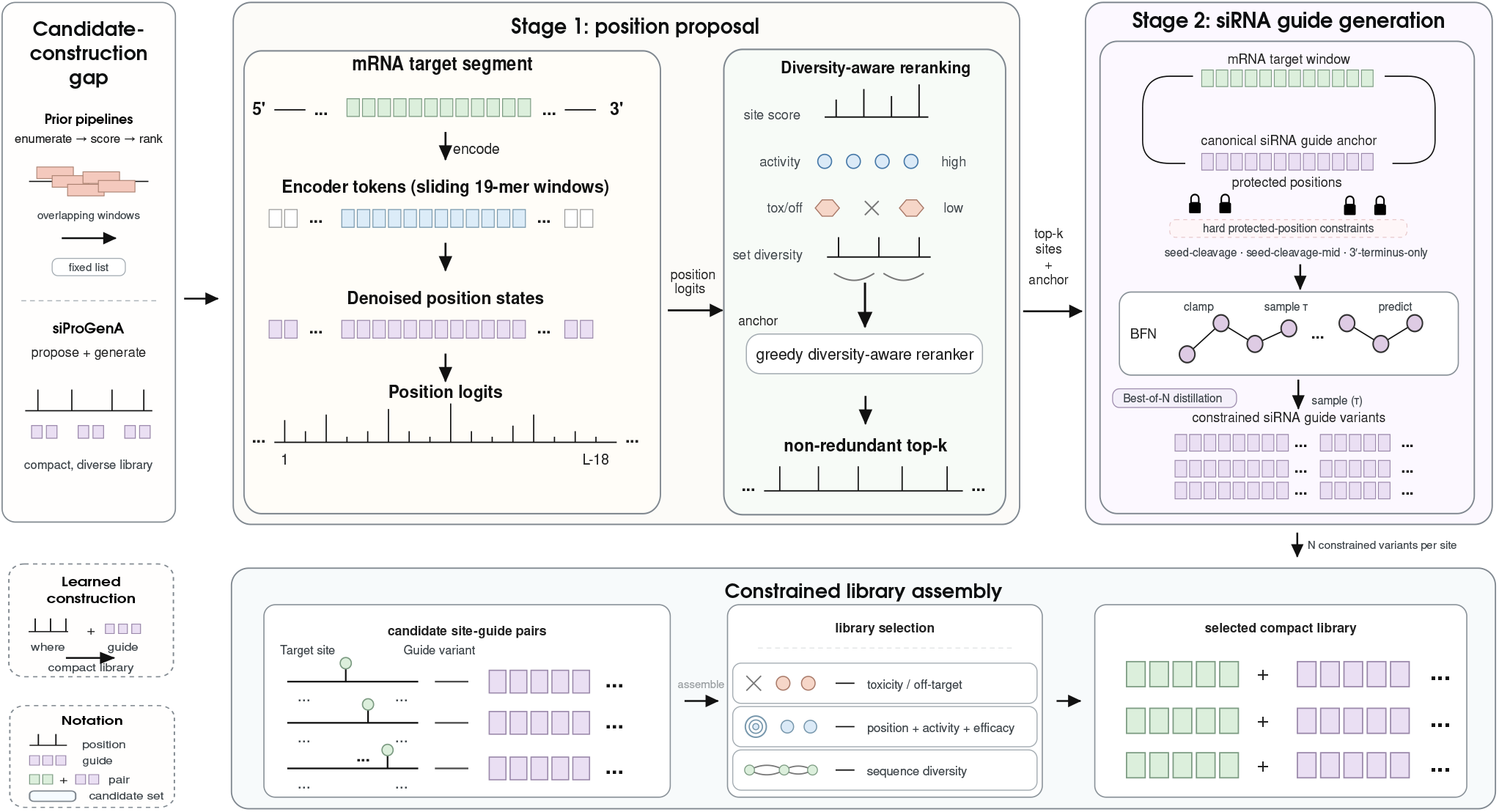
Overview of siProGenA. Stage 1 proposes a compact set of candidate target sites from an mRNA segment using mRNA-conditioned position proposal and diversity-aware reranking. Stage 2 generates *N* constrained siRNA guide variants for each selected site while preserving protected seed/cleavage positions. The constrained library assembly step then scores site-guide pairs under efficacy, toxicity/off-target, and sequence-diversity criteria to produce the final compact siRNA candidate library.

**Figure 2.**
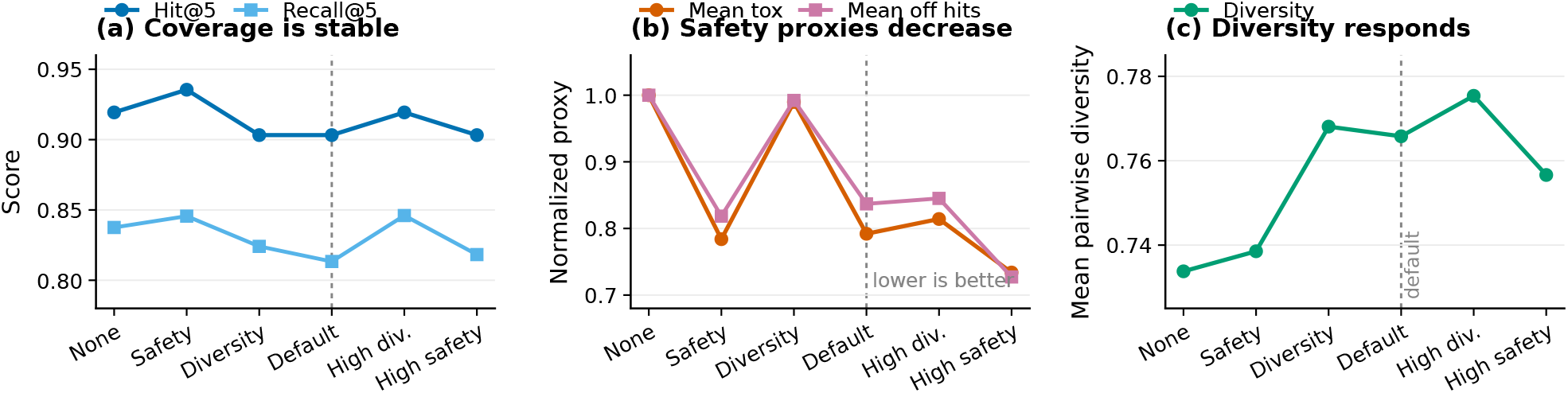
Reranking-weight sensitivity. Coverage remains stable across representative coverage–safety–diversity trade-offs. Safety-weighted settings reduce normalized toxicity and off-target proxies, while diversity-weighted settings increase pairwise guide diversity. The dashed line marks the default setting used in the main experiments.

### A.3 Additional Stage 2 Generation Results Figure 3 visualizes the paired comparison behind the main Stage 2 result. Most generated variants lie above the diagonal, showing that the learned constrained guide is often scored higher than the canonical complement at the same measured candidate position

Figure 4 shows that generated guides remain near the 0.45–0.55 GC design range rather than obtaining predictor gains through composition drift.

**Figure 3.**
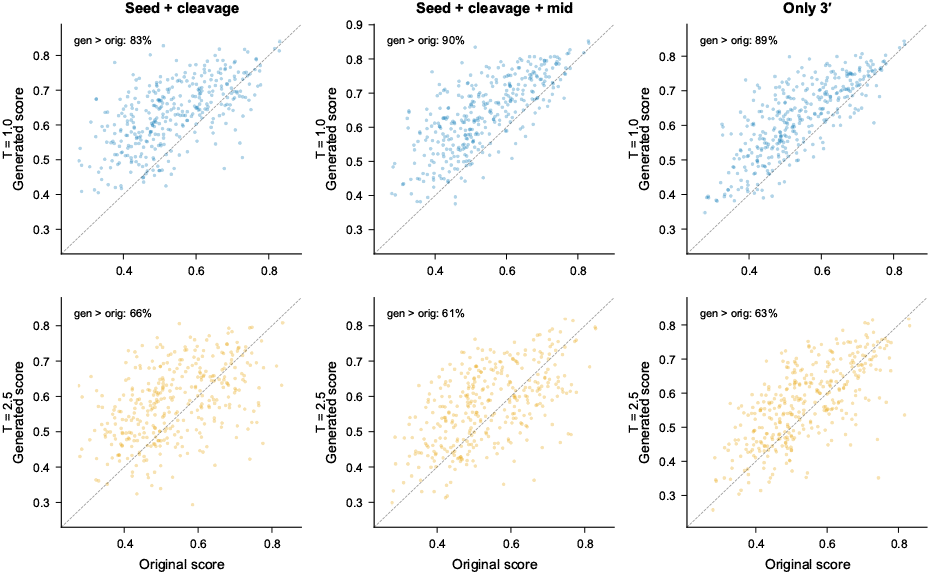
Constrained Stage 2 variants compared with the canonical complement under the external OligoFormer evaluator.

**Figure 4.**
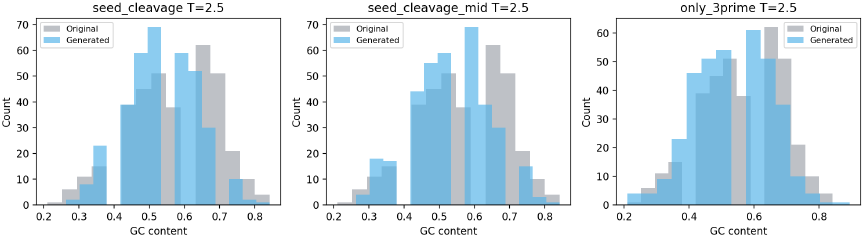
GC-content distributions for complementary guides and generated constrained variants at *T* = 2.5.

**Figure 5.**
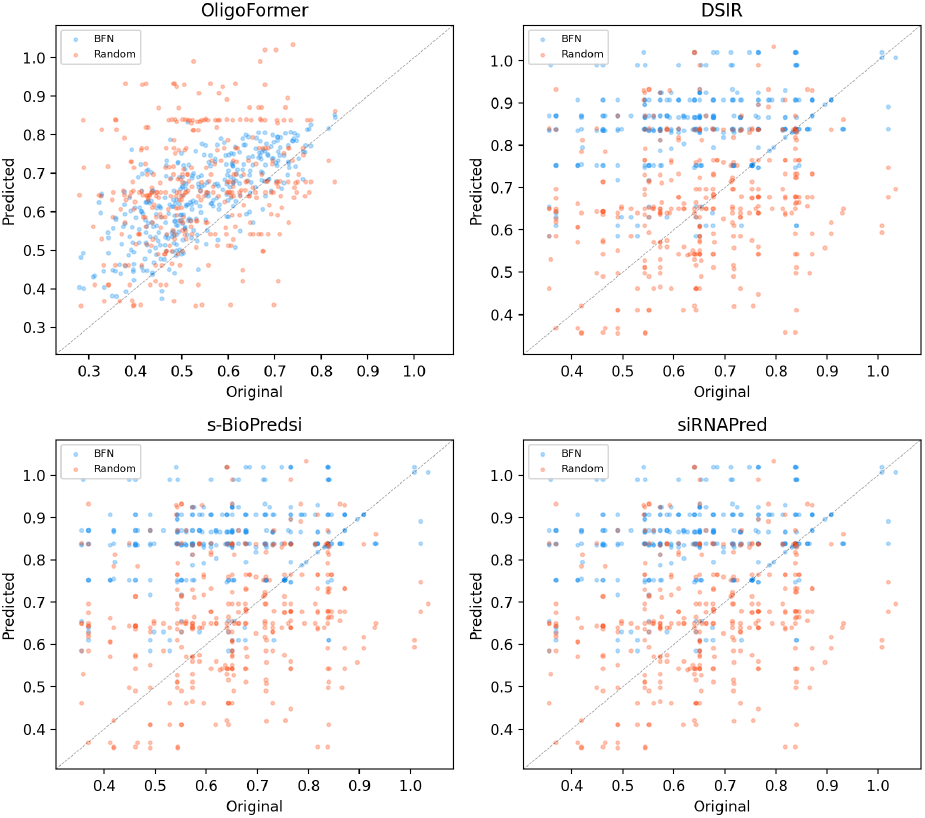
External-scorer comparison for learned BFN variants and random mismatch controls. Points above the diagonal indicate variants that outscore the canonical complement under the corresponding scorer.

Tables 10 and 11 expand the multi-model check across constraint modes and temperatures. The *T* = 1.0 setting gives the strongest mean-score and win-rate improvements, while *T* = 2.5 acts as a higher-diversity sensitivity setting with smaller but still positive mean gains.

**Table 10:** Four-model mean scores for constrained Stage 2 variants. Each model column reports original →generated mean score.

| Mode | $T$ | OligoFormer | DSIR | s-BioPredsi | siRNAPred |
| --- | --- | --- | --- | --- | --- |
| Seed+cleavage | 1.0 | 0.537→0.631 | 0.655→0.837 | 0.646→0.805 | 0.654→0.791 |
| Seed+cleav+mid | 1.0 | 0.537→0.635 | 0.655→0.842 | 0.646→0.834 | 0.654→0.737 |
| Only 3' variable | 1.0 | 0.538→0.618 | 0.655→0.848 | 0.646→0.821 | 0.654→0.726 |
| Seed+cleavage | 2.5 | 0.537→0.582 | 0.655→0.711 | 0.646→0.711 | 0.654→0.689 |
| Seed+cleav+mid | 2.5 | 0.537→0.570 | 0.655→0.723 | 0.646→0.716 | 0.654→0.692 |
| Only 3' variable | 2.5 | 0.538→0.568 | 0.655→0.719 | 0.646→0.704 | 0.654→0.682 |

**Table 11:** Four-model win rates for constrained Stage 2 variants.

| Mode | $T$ | Oligo | DSIR | sBio | siRNAP |
| --- | --- | --- | --- | --- | --- |
| Seed+cleavage | 1.0 | 82.8% | 84.7% | 85.3% | 87.9% |
| Seed+cleav+mid | 1.0 | 89.8% | 84.7% | 91.2% | 74.6% |
| Only 3' variable | 1.0 | 89.0% | 84.5% | 92.7% | 79.7% |
| Seed+cleavage | 2.5 | 65.5% | 58.8% | 62.1% | 61.0% |
| Seed+cleav+mid | 2.5 | 61.0% | 59.0% | 69.2% | 61.0% |
| Only 3' variable | 2.5 | 63.0% | 56.8% | 70.1% | 62.4% |

Table 12 reports the diversity-oriented *T* = 2.5 setting. Compared with the lower-temperature *T* = 1.0 setting in Table 5, the gains are smaller but remain positive across all constraint modes.

**Table 12:** Constrained guide generation at *T* = 2.5. This setting provides a diversity-oriented sensitivity check for Stage 2.

| Constraint mode | OligoFormer orig→gen | $\Delta$ | Win% | GC | Mismatch |
| --- | --- | --- | --- | --- | --- |
| Seed+cleavage | 0.537→0.582 | +0.044 | 65.5% | 0.535 | 8.20 |
| Seed+cleav+mid | 0.537→0.570 | +0.033 | 61.0% | 0.531 | 5.42 |
| Only 3' variable | 0.538→0.568 | +0.030 | 63.0% | 0.534 | 3.05 |

### A.4 Biophysical Validation of Constrained Guide Variants

We further evaluate the constrained guide variants with ViennaRNA 2.7.2 [22]. Table 13 reports differences from the canonical complementary guide at the same measured target site. Positive ΔMFE indicates weaker binding than the complement.

**Table 13:** ViennaRNA biophysical validation of constrained Stage 2 variants at *T* = 1.0 on 354 measured test sites. Values are differences from the complementary baseline. Smaller Δ duplex and seed MFE indicate better preservation of target binding; more negative terminal asymmetry favors guide loading.

| Sampling mode | $\Delta$ Duplex MFE | $\Delta$ Seed MFE | $\Delta$ Self-fold MFE | $\Delta$ Terminal asym. |
| --- | --- | --- | --- | --- |
| Seed and cleavage protected | +20.75 | +2.25 | +0.26 | -1.30 |
| Seed, cleavage, and mid-region protected | +14.72 | +2.24 | +0.47 | -1.30 |
| Only 3' terminus variable | +6.56 | +1.29 | +0.52 | -1.28 |

The constraint modes protect different guide positions. The seed-and-cleavage mode preserves complementarity at guide positions 2–7 and 10–11; the seed-cleavage-mid mode additionally protects the 3^′^ supplementary positions 13–16; and the 3^′^-terminus-only mode protects positions 1–15, allowing variation only at the guide 3^′^ terminus.

The four columns of Table 13 measure different biophysical properties. Duplex MFE measures the binding energy of the full guide– target duplex; a large positive Δduplex MFE means that mismatches substantially weaken on-target binding. Seed MFE measures binding in the seed region, where disruption is especially undesirable because seed pairing drives target recognition. Self-fold MFE measures whether the guide tends to fold onto itself; values closer to zero indicate weaker self-structure. Terminal asymmetry compares the stability of the two duplex ends, where more negative values favor guide-strand loading into RISC.

All values are relative to the canonical complementary guide. Protecting seed and cleavage-sensitive positions limits disruption in the recognition-critical seed region, while additional mid-region or 3^′^-terminus constraints further reduce full-duplex perturbation.

These results support the use of constrained guide variation: predicted-efficacy gains are obtained in a restricted sequence subspace rather than by arbitrary seed or cleavage-site disruption.

### A.5 Additional Random-Mismatch Control Figure 5 compares learned BFN variants and random mismatch variants across four external scoring models. The same generated and random candidates are plotted against the corresponding canonical complement scores. The learned variants show a more consistent upward shift than random mismatches, supporting the interpretation that Stage 2 learns a position-conditioned guide-variation rule rather than simply benefiting from non-complementarity

